# Deconvolution improves cryo-EM maps

**DOI:** 10.1101/2025.02.23.639707

**Authors:** Junrui Li, Wooyoung Choi, Yifei Chen, Shawn Zheng, Angus McDonald, John W. Sedat, David A. Agard, Yifan Cheng

**Author notes:** Correspondence and requests for materials should be addressed to Yifan Cheng. Correspondence. These authors contribute equally (Junrui Li, Wooyoung Choi).

## Abstract

With technological advancements in recent years, single particle cryogenic electron microscopy (cryo-EM) has become a major methodology for structural biology. Structure determination by single particle cryo-EM is premised on randomly orientated particles embedded in a thin layer of vitreous ice to resolve high-resolution structure in all directions. In practice, preferentially distributed particle orientations and/or other imperfections in imaging and data processing deteriorate quality of obtained cryo-EM map. Here we present a deconvolution approach, named AR-Decon, that computationally improves the quality of cryo-EM maps. We tested and validated the procedure, compared its performance with that of machine learning based density modification method, and benchmarked its performance with a wide range of deposited maps. Our results show that AR-Decon is robust and is a generally applicable post-processing procedure for single particle cryo-EM.

## Introduction

With the steady technological advancement over the past decade, single particle cryogenic electron microscopy (cryo-EM) has become a powerful tool for structure biology, enabling routine structure determination of many biological macromolecules at near-atomic resolution^1,2^. In this method, particles of the macromolecule being studied are randomly orientated in a thin layer of vitreous ice^3^. A three-dimensional (3D) density map is reconstructed from two-dimensional (2D) projection images of these particles, after selecting a homogeneous conformation by particle classification and assigning accurate Euler angles to each particle by iterative image alignment^4^. Given sufficient high-resolution projection images of randomly orientated particles, a 3D map can be reconstructed with an isotropic resolution. In practice, the resolution of a 3D map is often somewhat anisotropic^5,6^ and influenced by many factors, including orientational distribution of particles, image qualities characterized by signal-to-noise ratio (SNR) at both low- and high-frequency, and accuracies of image alignment, etc.

Mathematically, anisotropic resolution, or resolution in general, can be described by convolution, where experimentally obtained map is the ideal one distorted by convolving a point spread function (PSF) that describes the distortion of the map and attenuation of high-resolution SNR. A classic reverse computational approach is deconvolution^7^, which can in theory be applied to improve map quality by restoring the missing information and reducing attenuation of high-resolution SNR. Computationally, deconvolution algorithms treat the PSF as a kernel that acts on an object function, which is iteratively optimized by minimizing the differences between the experimental data and the object function convolved with PSF. The resulting object function approximates the ground truth data. *A priori* knowledge of the ground truth data is used to regularize the iterative processes. Deconvolution is extensively used in astronomy, spectroscopy, and imaging^8^. In recent years, it has also been applied to cryo-EM and cryo-STEM to fill the missing data in electron tomography^9-12^.

Most deconvolution algorithms developed for astronomy, spectroscopy or imaging are very sensitive to noise, severely compromising their applications when data is very noisy. This is particularly so for cryo-EM where the SNR of raw data is extremely poor. Entropy-regularized deconvolution (ER-Decon), developed by Arigovindan et al^13^, is a deconvolution program that was originally developed to restore images from widefield light microscopy with very poor SNR. In ER-Decon, the mathematic formulation of deconvolution is stated as

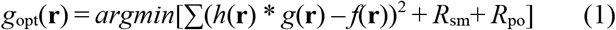

where **r** represents the pixel index, *g*(**r**) is the *predicted* ideal 3D widefield image that is to be optimized, *f*(**r**) is the measured widefield image, *h*(**r**) is the PSF. Two regularization functions, *R*_sm_ and *R*_po_, enforce smoothness and positivity. Different from many other deconvolution algorithms, such as traditional Landweber and Richardson–Lucy algorithms^14^, ER-Decon uses second-order derivatives in its regularization functions to suppress noise and enhance signal. Detailed explanations can be found in the original paper^13^ and its various applications^9-12^. Compared with other deconvolution algorithms, ER-Decon was documented to be able to handle much lower SNR^12^, making it more suitable for potential application in single particle cryo-EM reconstructions.

In this study, we apply ER-Decon to improve high-resolution SNR of cryo-EM maps, particularly those with anisotropic resolutions. Here, the above formula is re-written as

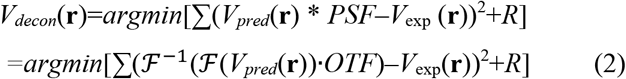

where ℱ and ℱ^−1^ denote forward and inverse Fourier transform, *V*_exp_ (**r**) is the final experimental map, *V*_*pred*_(**r**) is the *predicted* ideal map that is optimized during iterative process, OTF (optical transfer function) is the Fourier transform of PSF, and *R* represents regularizations. The deconvolution process takes an experimental map and its corresponding OTF as input and iteratively minimize the differences between the object map convolved with PSF and the experimental map. This iterative process optimizes the object map as closely as possible to the ideal map. Regularization combined with general real space constraints, such as positivity within region of interest, are generally critical for handling very low SNRs^9^.

The key step of applying ER-Decon is to generate an appropriate OTF from the target experimental 3D map and identify appropriate regularization parameters. We developed *AR-Decon* (AR-Decon: correcting <u>A</u>nisotropic <u>R</u>esolution by <u>Decon</u>volution), a procedure to generate an OTF from the two experimental half maps, followed by applying ER-Decon II algorithm to deconvolve the final experimental map. We validated the procedure with a test data generated from experimental data, optimized regularization parameters and benchmarked its performance by applying it to numerous experimental maps downloaded from EMDB. For maps with anisotropic resolution, AR-Decon improves the map quality by partially recovering the missing information in Fourier space. For maps with isotropic resolution, AR-Decon improves high-resolution SNR of the experimental map, enhancing high-resolution features without amplifying noise. We suggest using AR-Decon as a regular post-processing step to improve single particle cryo-EM reconstructions.

## Results

### Deconvolution of cryo-EM density map

An experimentally determined 3D cryo-EM density map can be considered as a perfect map convolved with a PSF and added with noise. The Fourier transform of an experimental map is then the product of the Fourier transform of the perfect map and the OTF. In ideal cases, where the angular distribution of all particles is uniform without any preferred orientation, the OTF would be spherical, isotropic in all directions, with an envelope function that describes the fall-off of SNR in Fourier components of the map towards high-resolution (Fig. 1a). Assuming a simple case where images within a certain angular range are completely missing, such as in random conical tilt^15^ or electron crystallographic reconstructions^16,17^, the OTF would be an incomplete sphere with an empty cone within which the information is missing (Fig. 1b). The PSF calculated from such an OTF with missing information is then distorted from the ideal spherical shape and the experimental density map calculated from such a dataset would be the ideal map convolved with the distorted PSF, causing elongation of the density map along the direction of missing cone (Fig. 1c).

**Figure 1.**
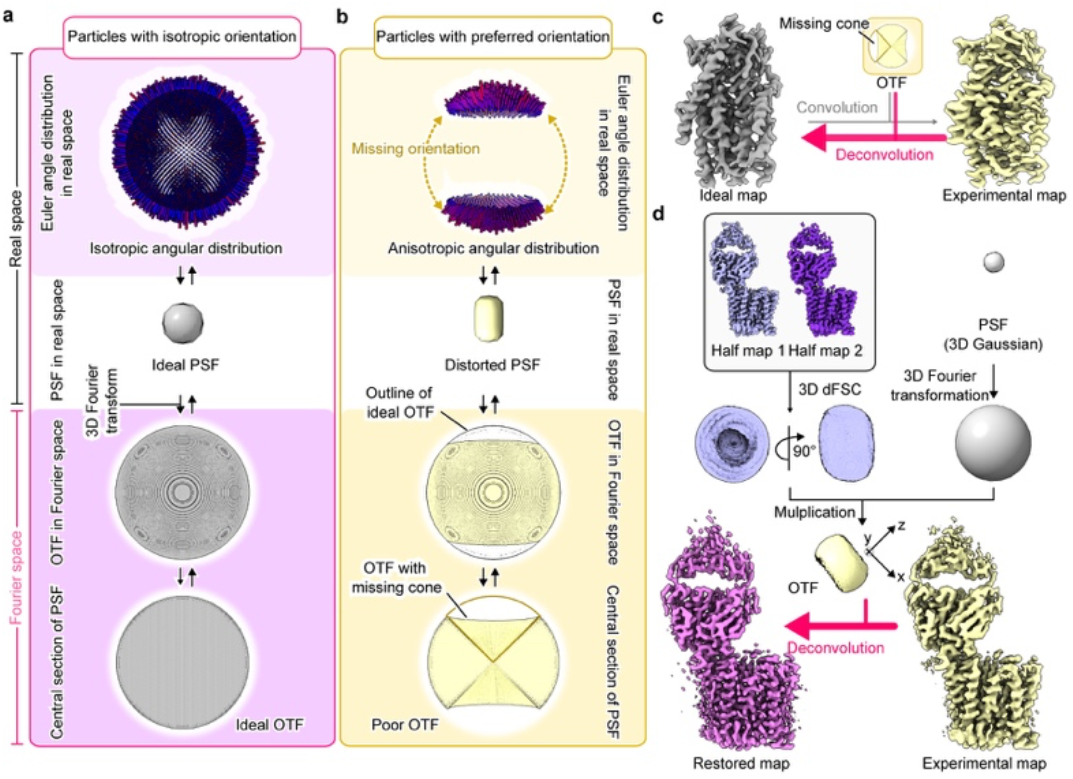
Principle and pipeline of deconvolution. **a**, Top to bottom, angular distribution of a dataset with evenly distributed particles, corresponding PSF in real space, and OTF in Fourier space, both are spheric. **b**, Top to bottom, angular distribution of a dataset with side views missing, corresponding elongated PSF, and OTF with a missing cone. **c**, An ideal map (grey) and a map with missing views (yellow) can be linked by convolution and deconvolution. **d**, In AR-Decon, OTF is generated from two experimental half maps, and is used to deconvolve the experimental full map to improve map quality.

In more general and practical cases, the angular distribution of particles is uneven with significantly fewer images along certain orientations often due to interaction with the air-water interface. The information distribution in Fourier space is then anisotropic, with poorer SNR and resolution in orientations with fewer particles. To generate an appropriate OTF, we first convert the directional Fourier Shell Correlation (dFSC)^18^ calculated from two experimental half-maps into a 3D map where the value of each voxel equals the corresponding FSC value. The OTF is then generated by multiplying this volume by the Fourier transform of a 3D spherical Gaussian PSF (essentially an overall B-factor to suppress buildup of high frequency noise) and then used for deconvolving the final experimental map to restore the map with improved and more isotropic resolution (Fig. 1d).

### Deconvolution partially recovers information in the missing cone

To validate and evaluate the performance of AR-Decon, we generated a test dataset from a high-quality experimental dataset collected on the ferroportin bound with a Fab^19^. The particles in the dataset are evenly distributed in Euler space, and a C1 reconstruction from this dataset has near isotropic resolution of ∼3Å. Fourier transform of the map shows no missing information in any direction. dFSC curves calculated from the two half-maps show very narrow distribution and the OTF generated from the dFSC is a sphere (Supplementary Fig. 1a-f). From this dataset, we generated a reconstruction with anisotropic resolutions by removing all particles with orientations more than 60° respect to the z axis, equivalent to a classic random conical tilt dataset with a 60° missing cone along the *z*-axis. Its Fourier transform shows a clear empty cone without information inside (Supplementary Fig. 1g-i). The overall resolution estimated from the FSC curve, which is equivalent to the average of dFSC along all directions, is only slightly worse than that of the original map. However, dFSC curves in different directions vary widely, with the resolutions along the directions inside the missing cone much worse than those outside (Supplementary Fig. 1j). The 3D representation of the dFSC and the corresponding OTF have a flat disk shape (Supplementary Fig. 1k, l). Note that calculating dFSC with a mask applied generates low resolution information in directions within the missing cone.

Deconvolution of the distorted map with its OTF using AR-Decon produced a “restored” map. Fourier transform shows that the empty cone is filled (Fig. 2a-c). To evaluate the quality of restored information (both amplitude and phase) within the missing cone, we calculated map-to-model dFSC of three maps, the original one, the one with missing cone, and the restored one (Fig. 2d-f). Map-to-model correlation within the missing cone is significant improved in the restored map, but, as expected, not to the level of the original map. The resolution difference between inside and outside the cone was significantly reduced. By the FSC=0.5 criterion, the average resolution estimated inside the cone was improved from 4.2Å (Fig. 2e) to 3.38Å (Fig. 2f) versus 3.17Å (Fig. 2d) in the original map.

**Figure 2.**
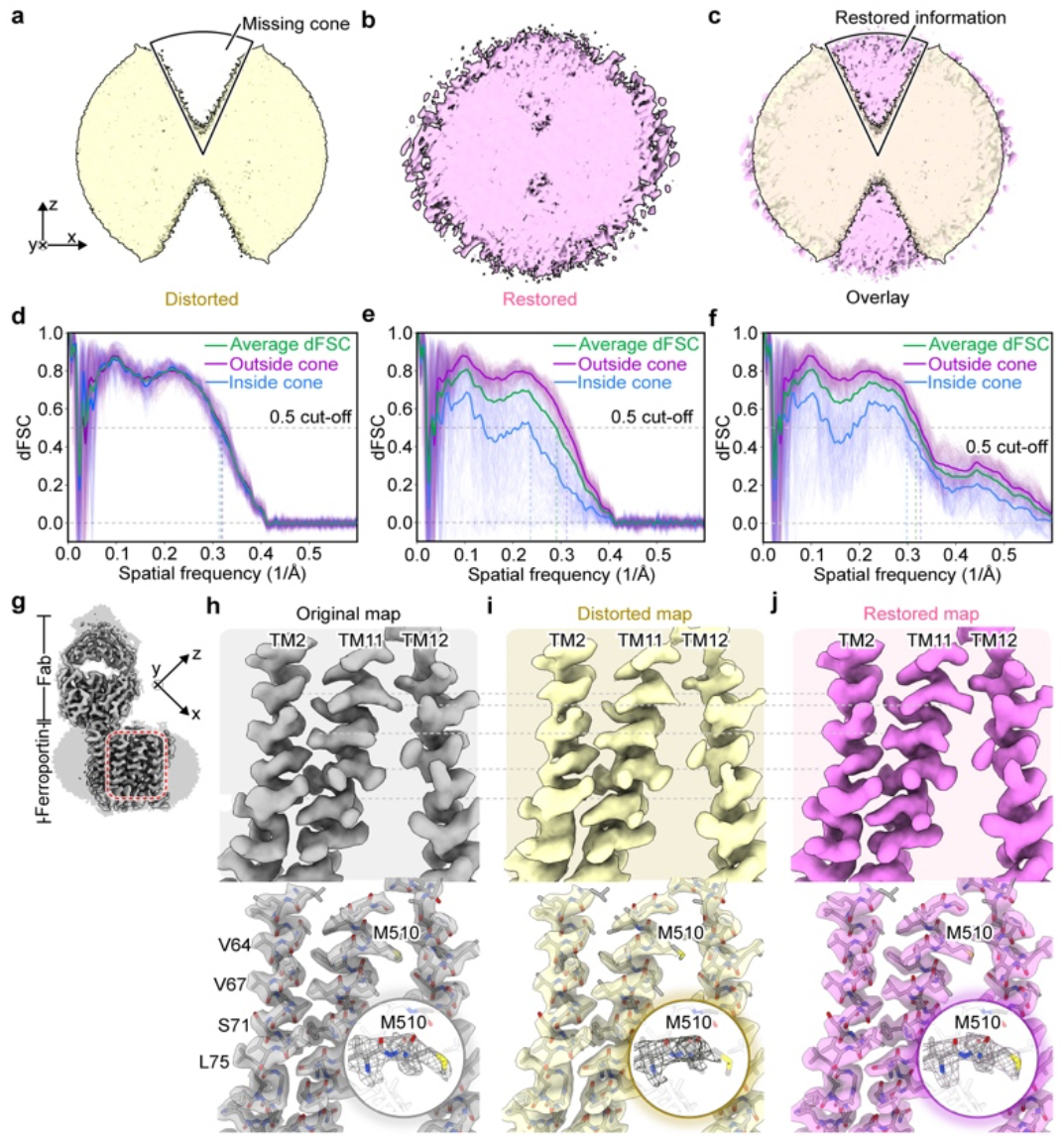
Deconvolution recovers partially missing information in Fourie space. **a**, 3D Fourier transform of distorted ferroportin-Fab map with a missing cone in z-direction. **b**, 3D Fourier transform of the restored map where the missing cone is filled. **c**, Overlay of (**a**) and (**b**). **d - f**, Map-to-model dFSC of the original (**d**), distorted (**e**), restored (**f**) maps. Thick lines indicate averaged dFSC curves of entire map (green), outside (purple) and within (blue) missing cone. **g**, Original ferroportin-Fab map. **h-j**, Enlarged view from boxed region in **g** of the original (**h**), distorted (**i**) and restored (**j**) maps. Upper row shows density of three transmembrane helices TM2, TM11 and TM12. Bottom row shows the same density with atomic model docked.

Deconvolution improves the distorted map. The elongations along the under-sampled directions are noticeably reduced. Focusing on a selected area of ferroportin (Fig. 2g), comparisons of the original (Fig. 2h) and distorted (Fig. 2i) maps show obvious differences, highlighting the influence of preferred orientations. Specifically, distorted map shows smeared and elongated density along the *z*-axis, which is particularly obvious for helix TM2. Side chain densities for residues V64, V67, S71, L75, M510 are smeared in the under-sampled directions (Fig. 2i). Deconvolution made noticeable improvement on the map by restoring side chain densities almost to the same level as the original map (Fig. 2j).

### Deconvolution of influenza hemagglutinin (HA) trimer

Single particle cryo-EM reconstruction of influenza hemagglutinin (HA) trimer (EMD-8731) reported an overall resolution of 4.2Å, with severe elongation caused by preferred orientation^20^. While β-strand and α-helices were resolved in the direction perpendicular to the preferred orientation, the map shows strong elongation along the symmetry axis. In this direction, the map is smeared to the extent that most of the side chain density is missing (Fig. 3a). In a later study, a structure of the same protein was determined to nearly isotropic 2.9Å resolution (EMD-21954, PDB 6wxb)^21^. We now use EMD-8731 to evaluate the performance of AR-Decon on a real experimental map and use EMD-21954 as the ground truth for comparison.

**Figure 3.**
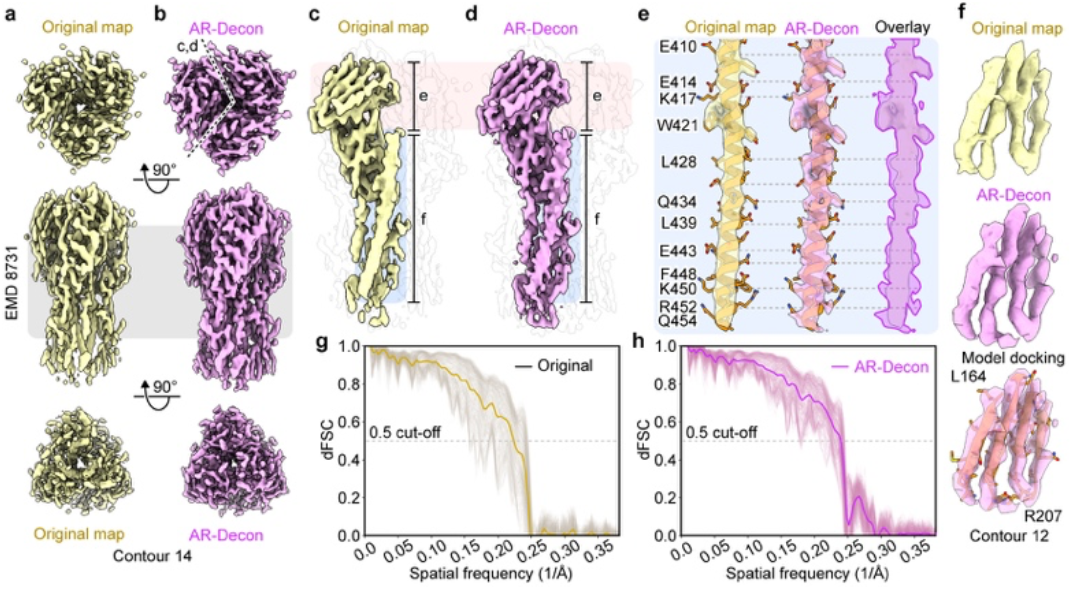
AR-Decon improves influenza hemagglutinin (HA) map. **a** and **b**, Top, side and bottom views of HA trimer map before (**a**) and after (**b**) deconvolution. **c** and **d**, Enlarged side view of a highlighted monomer before (**c**) and after (**d**) deconvolution. **e**, Density of an isolated α-helix from original (left) and deconvolved (middle) maps. Overlay of the two (right) highlights the improvement by AR-Decon. **f**, Density of an isolated β-sheet from original (left) and deconvolved (middle) maps. Docked atomic model (right) validates the modified densities (right). **g and h**, Map-to-map (EMD-21954) dFSC of the HA trimer map before (**g**) and after (**h**) deconvolution.

Following the same procedure, we generated an OTF from half map dFSC plots and used it to deconvolve the EMD-8731 map. After deconvolution, the smearing effect along the preferred orientation was noticeably reduced, with the helical turns and many side chains clearly resolved (Fig. 3b). A detailed comparison of local regions in the maps before and after deconvolution demonstrates the improvement from the deconvolution (Fig. 3c-f). For example, in the original map, β-strand densities were connected and helical turns were obscured. In the deconvolved map, connected β-strands were unambiguously separated and helical turns were clearly resolved, with side chain density restored to the level sufficient for model building. Using EMD-21954 as a reference map, we calculated the map-to-map dFSC for the maps before and after deconvolution (Fig. 3g, h). After deconvolution, the map-to-map dFSC exhibits reduced spreading, and the average dFSC also shows noticeable improvement.

### AR-Decon improves map resolution

In the ferroportin test described above, AR-Decon also improved the map-to-model dFSC outside the missing cone, from 3.21Å to 3.09Å (Fig. 2f). Likely, deconvolution would also improve the SNR of a target map even with isotropic resolution. To test this, we applied AR-Decon to the original deposited ferroportin map (EMD-21599). Indeed, AR-Decon improves its resolution, evaluated by map-to-model dFSC, from 2.82Å to 2.64Å (Supplementary Fig. 2). Visual inspection of the deconvolved density shows that it resembles B-factor sharpened map but with better connectivity in tracing backbone densities (Supplementary Fig. 2b, c).

To better understand the effects of AR-Decon and to differentiate it from B-factor sharpening, which scales the map Fourier amplitudes by an exponential function^22,23^, we performed a thorough test using another deposited map, human GAPDH (EMD-29664/PDB 8G17)^24^. It has a reported isotropic resolution of 1.98Å, and an estimated B-factor of -59.79Å^2^. Original and sharpened maps have almost identical dFSC curves, with the estimated resolution similar as the reported, using FSC=0.5 criterion. Applying AR-Decon to this map boosted the FSC curve with an improved resolution of 1.85Å, using the same resolution criterion (Fig. 4a). Thus, B-factor sharpening and deconvolution are different: sharpening only scales amplitudes without altering its SNR, but AR-Decon improves high resolution SNR. Considering that OTF envelop reflects the Fourier amplitudes of an experimental map, which is suppressed from the ideal map due to various factors, such as alignment errors or imperfections of raw micrographs, etc., our results suggest that the regularization strategy in the original ER-Decon algorithm help recover suppressed signals without amplifying noise.

**Figure 4.**
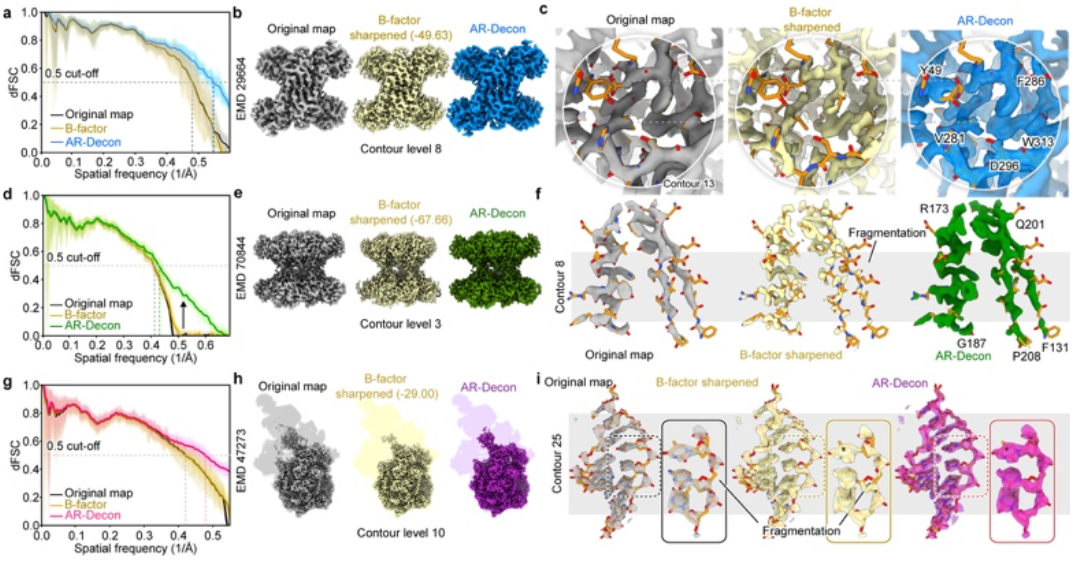
AR-Decon improves map resolution. **a, d** and **g**, Map-to-model dFSC of original (grey), B-factor sharpened (yellow) and deconvolved maps, as labeled. **b, e**, and **h**, Normalized original, B-factor sharpened and deconvolved maps (as labeled) are shown at the same contour level. Corresponding EMDB accession code are listed. **c, f** and **i**, Detailed comparison of specific regions of the original, sharpened and deconvolved maps. Sharpened maps are more fragmented than the deconvolved maps.

Deconvolved map appears similar as the sharpened map (Fig. 4b), in that holes of aromatic rings are visible in both sharpened and deconvolved maps. However, deconvolved map has noticeably better density connectivity, consistent with its improved high-resolution SNR (Fig. 4c, Supplementary Fig. 3a). Note that deconvolved map can be further sharpened by B-factor, but with a smaller B-factor of -41.15Å^2^, estimated using the same procedure.

We test AR-Decon with two additional maps (EMD/PDB: 70844/9OTP, and 47273/9BZ0) (Fig. 4d-i, Supplementary Fig. 3b, c). In both cases, B-factor sharpening does not alter the dFSC, whereas deconvolution improves the correlation between map and model in the high-resolution range. Visual inspection shows that deconvolved maps have better connectivity. For 47273/9BZ0, RNA densities are noticeably improved by deconvolution (Fig. 4i).

### Differences between AR-Decon and machine learning methods

Recently, several deep-learning based approaches, such as DeepEMhancer^25^, and IsoNet^26,27^, have been developed to improve the quality of maps having anisotropic resolution caused by preferred orientation. Another program, EMReady^28^, improves cryo-EM maps by modifying density in real space. Different from the explicit deconvolution method used in AR-Decon, which needs no training, deep-learning based methods rely on various training schemes. We explored their differences by applying AR-Decon and EMReady to two experimental maps, RNA Pol II elongation complex at the pre-catalysis state (EMD/PDB: 55239/9SV6)^29^ and GAPDH (29664/8G17)^24^.

The RNA Pol II elongation complex map has a reported resolution of ∼1.9Å, and contains many non-protein densities, including RNA, DNA and ATP surrounded by many water molecules^29^. EMReady improves the densities of the entire reconstruction rather impressively (Fig. 5a, b). However, it removes all densities interpreted as water molecules, likely being treated as noise. AR-Decon also improves protein density, although not to the same level as EMReady, but it preserves all non-protein densities, particularly all densities assigned as water molecules (Fig. 5c).

**Figure 5.**
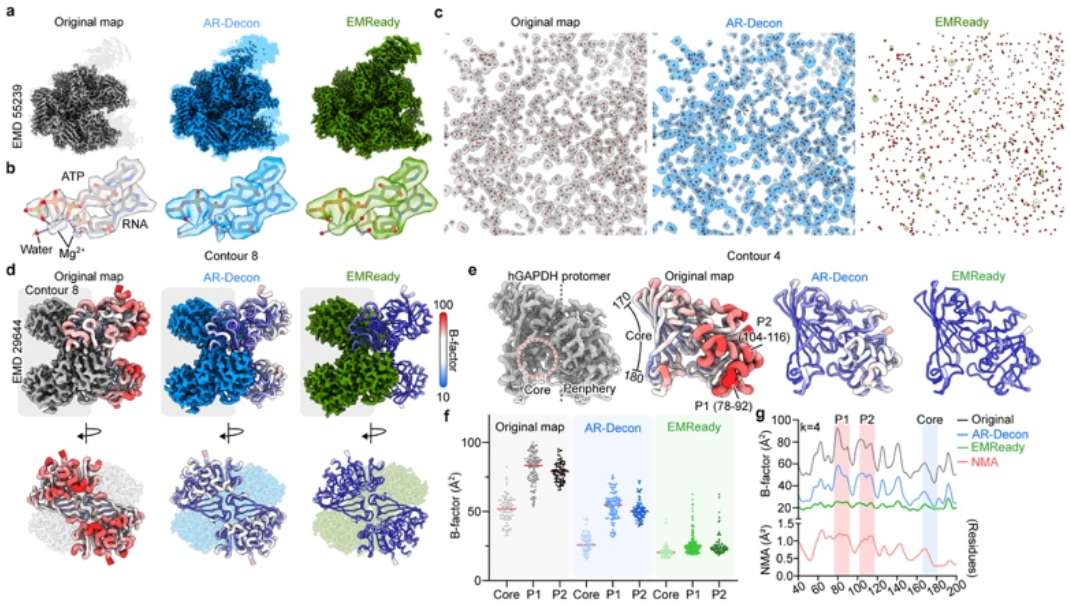
Comparison of AR-Decon and EMReady. **a**, Pol II elongation complex (EMD-55239): Original (grey), AR-Decon (blue) and EMReady (green) maps. All maps are normalized and shown at the same contour level. **b**, Densities of ATP, RNA, Mg^2+^ and water from three maps are compared. **c**, Water densities from each map are isolated with assigned water molecule (red dots). AR-Decon map preserves all water densities, while EMReady removes them. **d**, Two different views of human GAPDH (EMD-29644): Original (grey), AR-Decon (blue) and EMReady (green) maps of tetrameric GAPDH are shown together with the atomic model refined against each map. Atomic models are color coded by the B-factor assigned to individual residues during the refinement. **e**, Left panel: Density of a GAPDH protomer is isolated and shown; Second to fourth panels: Atomic model of protomer refined against original, AR-Decon and EMReady maps are shown in the same orientation, and color coded by the assigned B-factors, three regions, Core, Periphery 1 (P1) and Periphery 2 (P2), are labeled. **f**, Scatter plot of B-factors assigned to each residue in three regions of GAPDH protomer. AR-Decon map maintains the difference of B-factors assigned to different regions. EMReady map reduces B-factor in P1 and P2 regions, relative to the core. **g**, Plots of B-factor across the protomer, from residue 40 to 200, assigned to model of original (black), AR-Decon (blue) and EMReady (green) maps, and mobility calculated from the normal mode analysis. Shaded regions are labeled.

For GAPDH, EMReady improves the map density more impressively than AR-Decon, particularly in the peripheral region of the map. The root-mean-square-deviation (RMSD) between the atomic models refined separately against B-factor sharpened map, deconvolved map and EMReady treated map are ∼0.1Å, suggesting that the accuracies of both modified maps are reliable for atomic model refinement. However, the temperature factor assigned to each residue shows clear differences between these models (Fig. 5d-f). Dividing GAPDH protomer into three different subdomains, core (a.a. 170 ∼ 180), periphery 1 (a.a. 78 ∼ 92, P1) and periphery 2 (a.a. 104 ∼ 116, P2), B-factors of P1 and P2 regions are higher than that of the core in both original and deconvolved maps. In EMReady map, P1 and P2 have similar B-factors as in the core region. B-factors assignments in an atomic model refined against a specific density map are tightly correlated with the local resolution, which in turn reveal local conformational dynamics. In this case, the per residue mean square fluctuation calculated from normal mode analysis (NMA) of GAPDH correlate well with the B-factor assignment against the original and deconvolved maps but not with the EMReady map (Fig. 5g). Together, our analysis show that the original GAPDH map reveals that its periphery domains are more dynamic than the core. Such dynamic behavior is preserved in the deconvolved map, but not in the EMReady map, which could lead to the misinterpretation that the peripheral regions are as stable as the core.

### Using deconvolved maps as references improve angular refinement

3D map refinement in single particle cryo-EM is an iterative process in which the reconstruction from the previous round of refinement is used as the reference for the next iteration. In the case where particle distributions are somewhat preferred, reconstructions in the early stage of the refinement often show obvious elongation and their Fourier transform have missing information along the direction of the preferred orientation. Taking such a reconstruction as the reference for next round of angular refinement, it would be difficult to align particles oriented outside of the preferred orientation properly, since the correlation between such images and the reference would be weak. Thus, the preferred orientation problem would persist from the beginning, making it harder to recover experimentally when adding images that are outside of the preferred orientation. One idea to improve the situation is to repeatedly combine deconvolution and angular refinement, using the deconvolved map as the initial reference map in each refinement to improve the alignment of images oriented within the missing orientations (Supplementary Fig. 4a).

We tested this idea with an experimental dataset of TMEM16A, a calcium activated chloride channel (Supplementary Fig. 4b-f)^18^. Our original reconstruction from this sample shows clear preferred orientation. Here, we deconvolved the final reconstruction, used it as the reference and continued angular refinement (see Methods). The resultant map shows improvement, as the elongation of backbone density along vertical direction is reduced. Repeating the same procedure for one more round made further improvement of the reconstruction, as shown clearly by two-half map dFSC, with much narrower spread than the original reconstruction (Supplementary Fig. 4c-e). Similarly, the density, particularly side chain densities were noticeably improved (Supplementary Fig. 4f).

The same procedure of using deconvolved map as the reference map and continue angular refinement for more cycles could also be applied to datasets without obvious preferred orientations. As the refinement progresses, using the deconvolved map with improved high-resolution SNR as the reference could improve further alignment accuracy.

### Benchmarking AR-Decon

To test the broad applicability of AR-Decon, we evaluated its performance by applying it to various maps selected from EMDB database, divided into two categories: maps with anisotropic resolutions, and those having isotropic resolutions.

For the first category, we selected following maps (EMD) and coordinates (PDB): E41062/8T60, 41138/6TAN, 42494/8URJ, 50018/9EVW, 62392/9KKM. For each entry, we downloaded two half maps and the final map without sharpening for deconvolution. The results were evaluated by visual inspection and by map-to-model dFSC (Supplementary Fig. 5). In all these maps, AR-Decon improved map quality by reducing its anisotropy to various degrees. Using the deconvolved map as a reference for further particle refinement may improve the map further, but we leave this to be tested by the owners of these maps.

We also tested the following maps without obvious anisotropic resolution: 71540/9PDQ, 47016/9DMU, 60326/8ZP4, 70222/9O8C, 70660/9OO6, 49386/9NGJ, 29873/8G9O and 47658/9E75 (Supplementary Fig. 6 and 7). All these maps contain non-protein densities, such as lipids, ions, small molecules, or nucleic acids. AR-Decon made obvious improvements in all maps, with non-protein densities better resolved. One specific example is 49386/9NGJ, which is an ABC transporter with phytanoyl-CoA bound (Supplementary Fig. 6g). AR-Decon improves the shape of phytanoyl-CoA density in this map. In comparison, EMReady produces better improved protein densities but removes phytanoyl-CoA density.

Overall, these extensive tests show that the performance of AR-Decon is target independent and can be applied broadly to almost any type of reconstruction.

## Discussion

In this study, we established a workflow, AR-Decon, that applies the ER-Decon deconvolution algorithm to improve map quality for single particle cryo-EM. We first derived an OTF from dFSC plots between two half-maps, followed by using ER-Decon II to deconvolve this OTF from the experimental final map. For maps with anisotropic resolutions caused by preferred particle orientations combined with misalignment, AR-Decon improves information recovery along the poorly resolved directions in Fourier space. In many cases, incorporating AR-Decon into refinement further improves the anisotropic resolution of the final map. For maps with isotropic resolution, AR-Decon enhances high-resolution features. Benchmarking tests show that all maps processed by AR-Decon have some level of improvement, demonstrating the broad applicability of AR-Decon.

While related, constrained and regularized deconvolution and B-factor sharpening are fundamentally different. Due to these additional features, AR-Decon improves the high-resolution SNR and map-to-model correlation by recovering signals suppressed by the envelope function that reflects the combined effects of preferred particle orientation, imperfections in raw data acquisition and processing. B-factor sharpening scales the map amplitude to enhance high-resolution feature^22,23^, but does not change the SNR or map-to-model correlation. Over sharpening by the exponential function makes the map noisier and thus can fragment connectivity. Thus, instead of using B-factor sharpening as a standard postprocessing step, we suggest applying AR-Decon first, followed by further sharpening with a smaller B-factor. This often generates a better map, as demonstrated in this study.

It is worth noting the difference between AR-Decon and machine learning based density modification methods. Machine learning based methods^25-28^ use pre-trained generative AI model to modify map densities in real space directly. Limited by the training scheme and data used for training, these methods have the potential to over correct densities, such as removing real but non-protein densities or reducing local conformational dynamics captured in maps. As a pure mathematic procedure with tunable regularization parameters, although non-linear, AR-Decon is target independent and does not hallucinate features. Due to its advanced regularization, it is a stable procedure that does well to correct for anisotropic resolution as well as general high-resolution detail without amplifying noise. It does not reduce but rather preserves variations of local resolution within a reconstruction, which are typically caused by local conformational heterogeneity within particles or protein dynamics.

There are other mathematic based approaches, such as ResolveCryoEM implemented in Phenix^30,31^, which modify density by enforcing solvent flattening and histogram matching in combination with maximum likelihood recombination in Fourier space^30,31^. We have not tested ResolveCryoEM extensively to compare its performance with AR-Decon. Nevertheless, it is important to keep in mind that, in practice, different methods may produce somewhat different results in any given situation.

We recognize that the deconvolution methods require some estimate of the true OTF. Our strategy to estimate the OTF from the dFSC may miss some details as it is calculated by rotating a cone with a 40º apex angle. While this captures the overall behavior well, it would be interesting to explore other approaches, such as deriving the OTF from the D_obs_ value used in the EM_placement procedure, which provides a per-Fourier-term estimates of the local FSC in Fourier space^30^.

Since the conventional method of estimating resolution from FSC of two experimental half maps is no longer available for deconvolved maps, final resolution assessment of the deconvolved maps has to be evaluated by another method. In this study, we use dFSC between the deconvolved map with the atomic model. Thus, we currently can only evaluate the final deconvolved map quantitatively if a reliable atomic model is already available. Otherwise, visual inspection must be used to compare the deconvolved map, providing a qualitative evaluation of the map improvement.

## Methods

### Description of AR-Decon

AR-Decon package contains following parts: generating OTF, applying ER-Decon II^13^ to deconvolve OTF from the target map, optional screening of regularization parameters to further optimize performance of ER-Decon. Furthermore, it uses map-to-model dFSC to evaluate the deconvolved map, and bfactor.exe to further sharpen the deconvolved map by a negative temperature factor.

### Directional Fourier shell correlation (dFSC)

dFSC was described previously^18,32^. Briefly, it takes a pair of unfiltered and unsharpened half maps as input. Instead of calculating correlation between two half maps at each spherical shell in Fourier space, the correlations within conical shells in different directions are calculated. We evenly sampled 500 directions using Fibonacci approach. For each direction, all the voxels within the cone centered along this direction with apex angle of 40° were involved in the calculation. Along each direction and within the cone, a Fourier shell correlation is calculated. Once all 500 dFSCs were calculated, a 3D dFSC volume is rendered by calculating weighted average for each voxel. A script of calculating dFSC is included in the package. To calculate map-to-model dFSC, an atomic model is first converted to map by using either UCSF Chimera^33^ or ChimeraX^34^ with *molmap* command.

### Optical transfer function (OTF)

OTF is constructed from two experimental half maps (Fig. 1d). We first calculate dFSC from two half-maps, followed by generating a volume representation of dFSC. Briefly, a 3D gaussian function

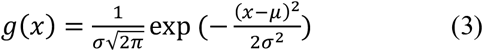

with its parameters (μ,σ) initialized as below:

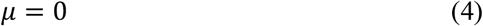

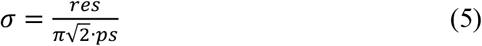

is generated, where *res* is the resolution estimated by half-map FSC at threshold 0.143 and *ps* is the pixel size of the map. Equation (3) makes FT of the gaussian distribution fall to 1/*e* of its maximum value at frequency 1/*res* , following the definition used *molmap* command in UCSF Chimera. OTF is constructed by multiplying Fourier transform of this Gaussian function by the 3D representation of dFSC.

### Mask

A mask for the target map can be used for calculating dFSC and for applying AR-Decon. Spheric or ellipsoid mask with soft edge are commonly used mask. To generate such a mask, the final total map is displayed with a proper threshold in UCSF Chimera or ChimeraX to identify the boundary and the dimension of the map. A soft spheric or ellipsoid mask can be generated using either UCSF Chimera or ChimeraX. A script of generating spheric mask is included in the AR-Decon package. This mask is then applied to two half-maps, before calculating dFSC, or as an input for deconvolution.

### Pipeline of applying AR-Decon

The pipeline of applying AR-Decon is described in Supplementary Fig. 8a. It starts with running AR-Decon with default setting, iterations = 50, smoothing = 0.5 and nonlinearity = 10,000. This is followed by optional procedure to optimize regularization parameters and to test longer iteration number. Although unnecessary in most cases, the optimization procedure can be repeated multiple times if desired. With the current version of ER-Decon II, which runs on a single CPU core, computation time of each deconvolution run is directly correlated with number of iterations, which imposes a practical limitation to performing extensive optimization. Efficient optimization would require distributing the process across multiple CPU cores.

AR-Decon requires two half maps and a full map as inputs. The full map can be either unsharpened or sharpened by a negative B-factor. While optional, providing a mask often yields better result. Note that AR-Decon should not be applied to two half maps, because deconvolving two independent half maps with the same OTF yields two maps that are no longer independent from each other. However, without two independent half maps, resolution and B-factor of deconvolved map cannot be estimated by the standard procedure, and auto-sharpening procedure implemented in Relion or Phenix cannot be applied. Instead, the resolution of deconvolved map is estimated by map-to-model dFSC and bfactor.exe (https://grigoriefflab.umassmed.edu/bfactor) is used to estimate the B-factor and to sharpen the deconvolved map.

### Iteration number and regularization parameters

ER-Decon II algorithm iteratively minimizes the loss function in formula (2) to produce an outcome that is as close as possible to the ground truth. Prolonged iterations produce sharper features in the deconvolved map, resembling more to the effect of over-sharpening by B-factor (Supplementary Fig. 8b). Two regularization parameters are smoothing that strengthens real-space smoothness to suppresses high-frequency random noise, and nonlinearity that preserves weak high-resolution signal by enforcing sparsity (Supplementary Fig. 8c). Therefore, interplays of iteration number, smoothing and nonlinearity ensure that AR-Decon recovers true high-resolution signals suppressed by imperfections in data or data processing without amplifying noise.

### Generating and processing ferroportin test dataset

The final dataset that produced deposited ferroportin^19^ reconstruction contains 310k particles. Instead of using the original deposited map determined from this dataset, which was further processed by polishing, we used Euler angles assigned during the refinement and calculated a reconstruction at an isotropic resolution of 3.1Å and used it the original (ground truth) map in our test. For each particle in this dataset, we derived a direction vector from refined Euler angles and then calculate angle between its directional vector and the z-axis. A subset of particles was generated by removing all particles with the angle outside the range of 0° ∼ 60°, or 120° ∼ 180°. The resulting subset contains about 178k particles, and their angular distribution was calculated using star2build.py in pyem (https://github.com/asarnow/pyem) and displayed using UCSF Chimera. The angular distribution verified that the preferred orientation was indeed along *z* -axis (Supplementary Fig. 1h). A reconstruction calculated from this subset has a missing cone in its Fourier space and is used as distorted map. Two half-maps were also calculated to generate OTF for deconvolution. Both original and distorted maps were sharpened using phenix.auto_sharpen tool^23^, with B-factor of -54Å^2^ and -37Å^2^, respectively. A spheric mask was generated for the distorted map at a contour level 0.02, with edge width of 20 pixels. Deconvolution was performed on the distorted full map following the pipeline described above with a spheric mask. The deconvolved map was normalized using the “normalize” processor of e2proc3d.py in EMAN2^35^.

All the maps shown in Fig. 2h-j were displayed at the same contour level of 8.5 using UCSF ChimeraX.

### Optimization of iteration and regularization parameters

Using ferroportin with a missing cone as the test data and map-to-model dFSC within the missing cone of deconvolved map as the screening criterion, we optimized iteration, smoothing and nonlinearity. The parameters obtained, 50 iterations, smoothing = 0.5 and nonlinearity = 10,000, are found to produce results that are sufficiently good in all maps we tested (Supplementary Fig. 9). Thus, they are used as default settings in the AR-Decon pipeline. A script is included in the AR-Decon package to enable further optimization of smoothing and nonlinearity for any target map. It outputs a gallery of dFSC panels for convenient evaluating the result.

An efficient way to optimize the performance is to start AR-Decon with default settings, followed by a grid search of the two regularization parameters, and identifying new parameters from the output gallery of dFSC. Using the optimized regularization parameters, more iteration cycles can be further tested. If a longer iterative cycle is to be used, it is advisable to screen for regularization parameters again. While the optimization procedure can be repeated multiple times, our experience shows that more than one round of optimization is unnecessary. The ranges of grid search used to identify the default settings are that smoothing: 5e-5, 5e-4, 5e-3, 1e-2, 2e-2, 5e-2, 1e-1, 2e-1, 5e-1, 1, 2, 5, 1e1, 2e1, 5e1, 1e2, 2e2 and 5e2, nonlinearity: 1, 1e1, 1e2, 1e3, 1e4, 1e5, 1e6 and 1e7, and iteration: 5, 20, 50, 100, 500, 1000, 2000, and 5000. An example of further parameter search of GAPDH (EMD-26944) is shown in Supplementary Fig. 10. The final parameters used for deconvolving the experimental map is marked by a star.

### Influenza hemagglutinin (HA) trimer

The HA trimer reconstruction, two half maps and the mask were downloaded from EMDB under accession number 8731^20^. dFSC between two half maps was calculated with the deposited mask, and a corresponding OTF was generated. (Note that the deposited two half-maps and the mask of EMD-8731 have the opposite handedness and need to be first flipped before further processing.) AR-Decon was applied to the post-processed map with default settings. The deconvolved map was sharpened using the auto sharpening tool^23^ in Phenix^36^ and the nominal resolution was set at 4.2Å. To calculate dFSC between deconvolved map with EMD-21954, two maps need to be first aligned and resampled to have the same pixel size and box dimensions using UCSF ChimeraX.

### Incorporating AR-Decon into 3D refinement

Repeated refinement and deconvolution were performed on the TMEM16A dataset. In the first round of processing, the TMEM16A particle stack was refined using non-uniform refinement^37^ tool in CryoSPARC^38^ and yielded a full map and two half maps. The refined full map was deconvolved with the OTF and the default smoothing and nonlinearity parameters. In the second round of processing, the deconvolved map was low pass filtered to 6Å and served as an initial reference for the second round of non-uniform refinement, followed by the second round of deconvolution. Finally, the third round of processing was performed in the same way as the second one.

### Normal Mode Analysis

Normal Mode Analysis (NMA) of human GAPDH (PDB:8g17) was performed using IMODs (https://imods.iqf.csic.es/)^39^ with default settings. For Fig. 5g, extracted B-factor from each atomic models and mobility from NMA was plotted using Prism Graphpad 10. For better visualization, rolling average method was applied (k=4, polynomial 2^nd^ orders).

## Data Availability

Deconvolved map from the original deposited map of influenza hemagglutinin (HA) trimer (EMD-8731) is deposited to EMDB with access code EMD-49155.

## Code availability

AR-Decon that contains python scripts to generate OTF for deconvolution and the executable of ER-Decon II is released in GitHub as open source (https://github.com/yifancheng-ucsf/AR-Decon).

## ACKNOWLEDGMENTS

We are grateful to Eric Branlund for the instruction of using ER-Decon. We thank J. Zhao, S. Li, and D. Asarnow for discussion, C.M. Azumaya, S. Dang, S. Feng and C. Puchades, B. Faust for providing their dataset for testing. This work was supported by grants from the National Institute of Health (R35GM140847, and U54AI170792) to Y.C. The UCSF cryo-EM facility was partially supported by NIH grants (S10OD020054, S10OD021741, and S10OD025881). Y.C. is an investigator of the Howard Hughes Medical Institute.

## AUTHOR CONTRIBUTIONS

J.W.S. suggested the initial idea of applying ER-Decon to process cryo-EM datasets. J.L. and Y. Cheng. designed the study. J.L., J.W.S., D.A.A and Y. Cheng designed deconvolution pipeline. J.L. implemented the deconvolution procedure and carried out related studies. W.C. performed evaluation and tests of applying AR-Decon to improve isotropic resolution and SNR and performed benchmarking tests. Y. Chen conducted 3D refinement using the deconvolved map as references. S.Z., J.W.S. and D.A.A. provided insight and suggestions throughout the study. A.MD. updated and maintained ER-Decon program. J.L., W.C. and Y. Cheng. wrote the manuscript with input from all authors.

---

**Supplementary Figure 1.**
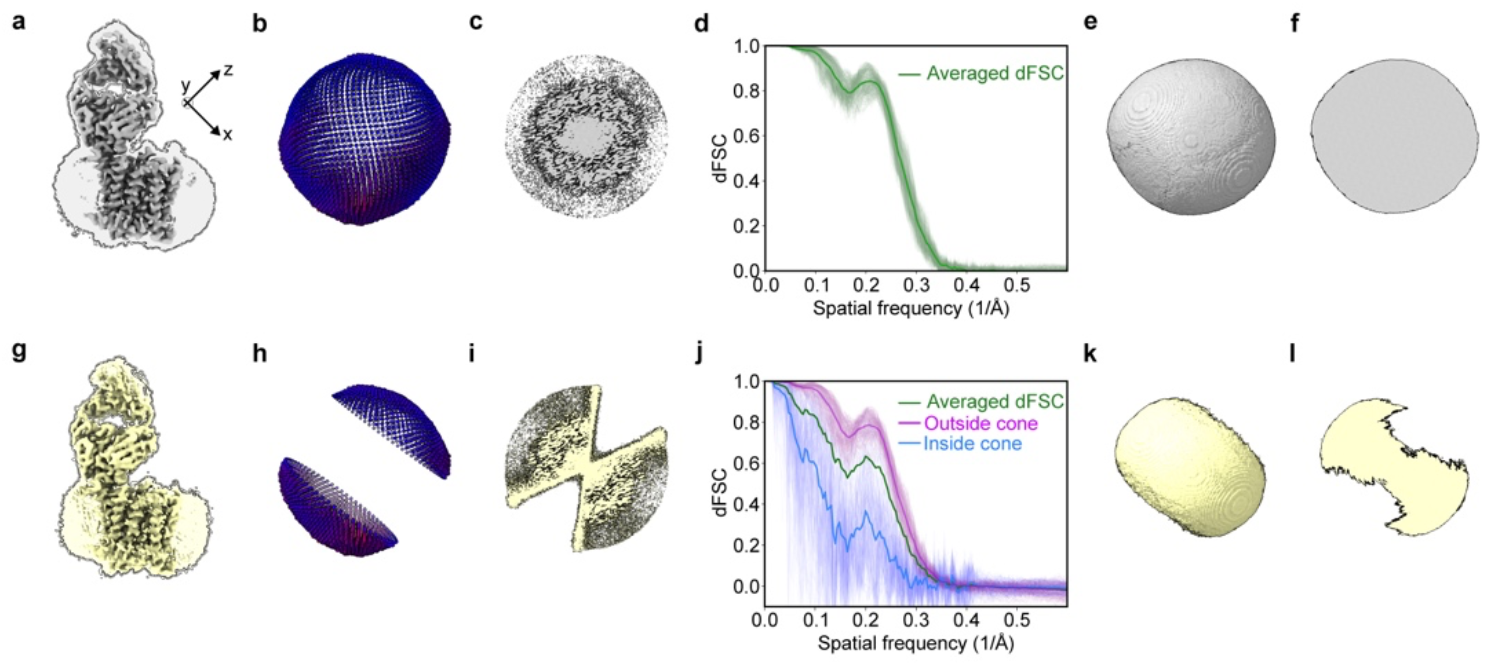
Generating ferroportin-Fab reconstruction with a missing cone. **a**, Reconstruction of ferroportin-Fab complex determined a dataset with uniform angular distribution. **b**, Angular distribution of the dataset. **c**, Central z-slices through 3D Fourier transforms of map shown in (**a**). **d**, dFSC curves calculated between two independent half-maps. The thick green line represents the average dFSC, while the thin green lines show individual dFSC curves for all directions. **e** and **f**, 3D dFSC volume (**e**) rendered from (**d**) and its central z-slice along z-axis (**f**), demonstrating isotropic resolution. **g**, Distorted reconstruction obtained from the dataset in which all particles with orientations more than 60° with respect to the z axis are removed. **h**, Angular distribution of z-axis as the preferred orientation. **i**, Central z-slices through 3D Fourier transforms of the distorted map. **j**, dFSC curves calculated between two independent half-maps refined from particles with z-axis preferred orientation. Thin blue and purple lines are dFSC curves inside and outside the cones, respectively. **k** and **l**, 3D dFSC volume (**k**) derived from (**j**) and its z-slice (**l**), revealing missing information and reduced resolution along z-axis, the axis of preferred orientation.

**Supplementary Figure 2.**
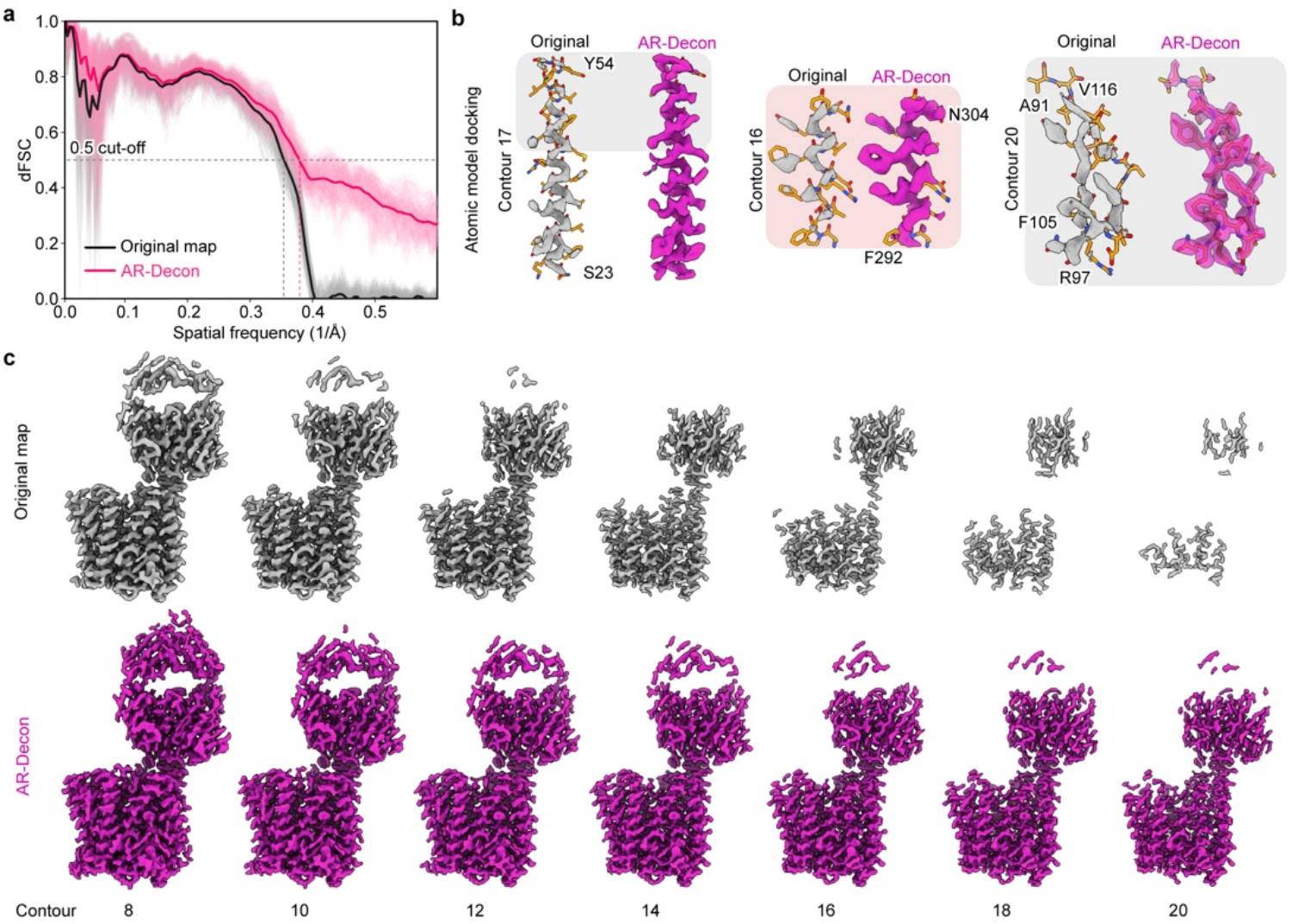
Apply AR-Decon to EMD-21599. **a**, Map-to-model dFSC of original deposited ferroportin-Fab map (EMD-21599, black curve) and deconvolved map (red curve). FSC=0.5 criterion is marked. **b**, Selected regions of the original (gray, left) and deconvolved map (purple, right) are displayed at the same contour level. **c**, Original (grep, upper panel) and deconvolved (purple, lower panel) are displayed at different contour levels. Deconvolved map has better density connectivity.

**Supplementary Figure 3.**
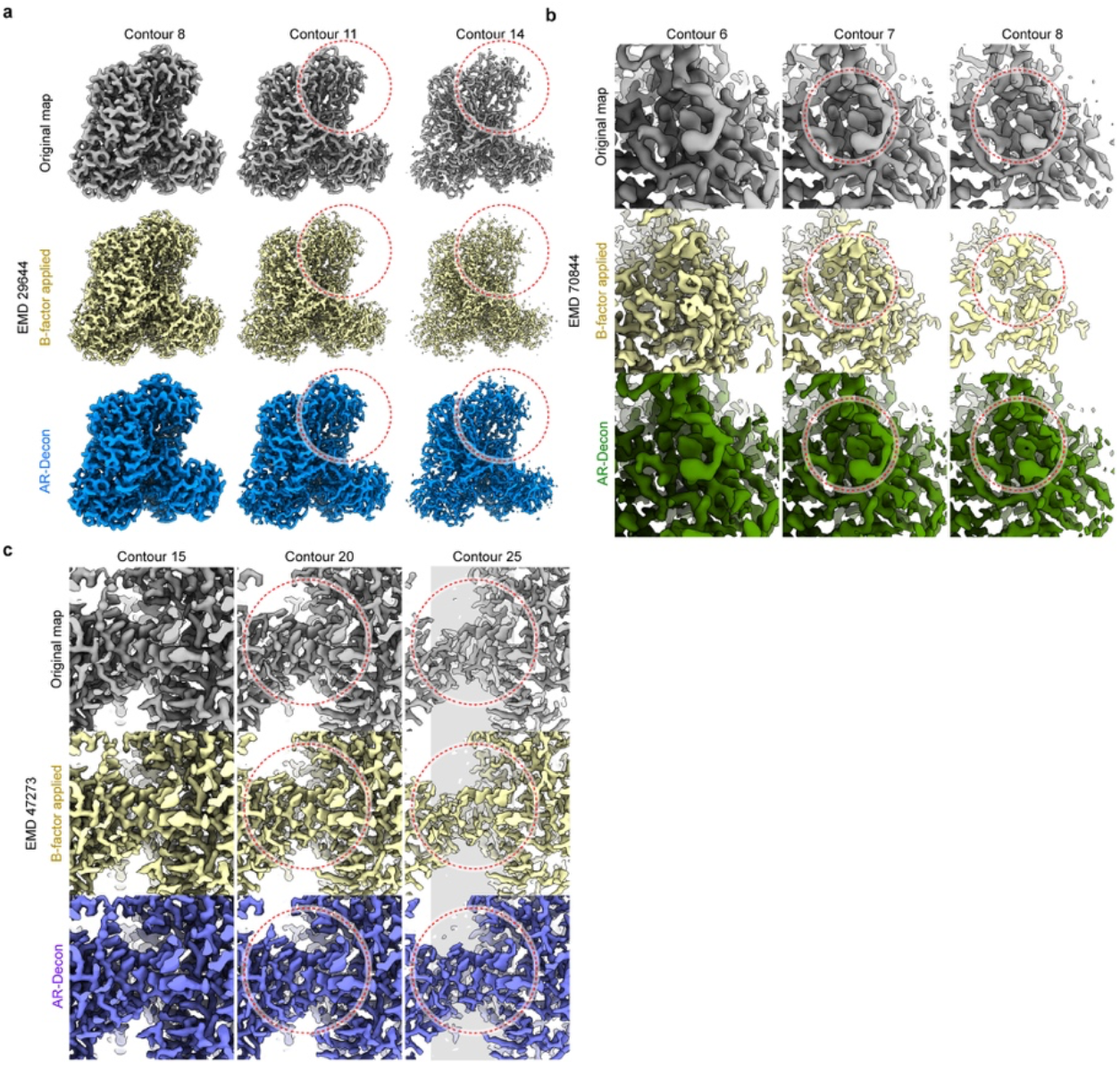
Comparison of B-factor sharpening with deconvolution. **a, b**, and **c**, Original (grey), B-factor sharpened (yellow) and deconvolved maps (EMD-29644 (**a**), -70844 (**b**) and -47273 (**c**)) are compared at different map contour levels.

**Supplementary Figure 4.**
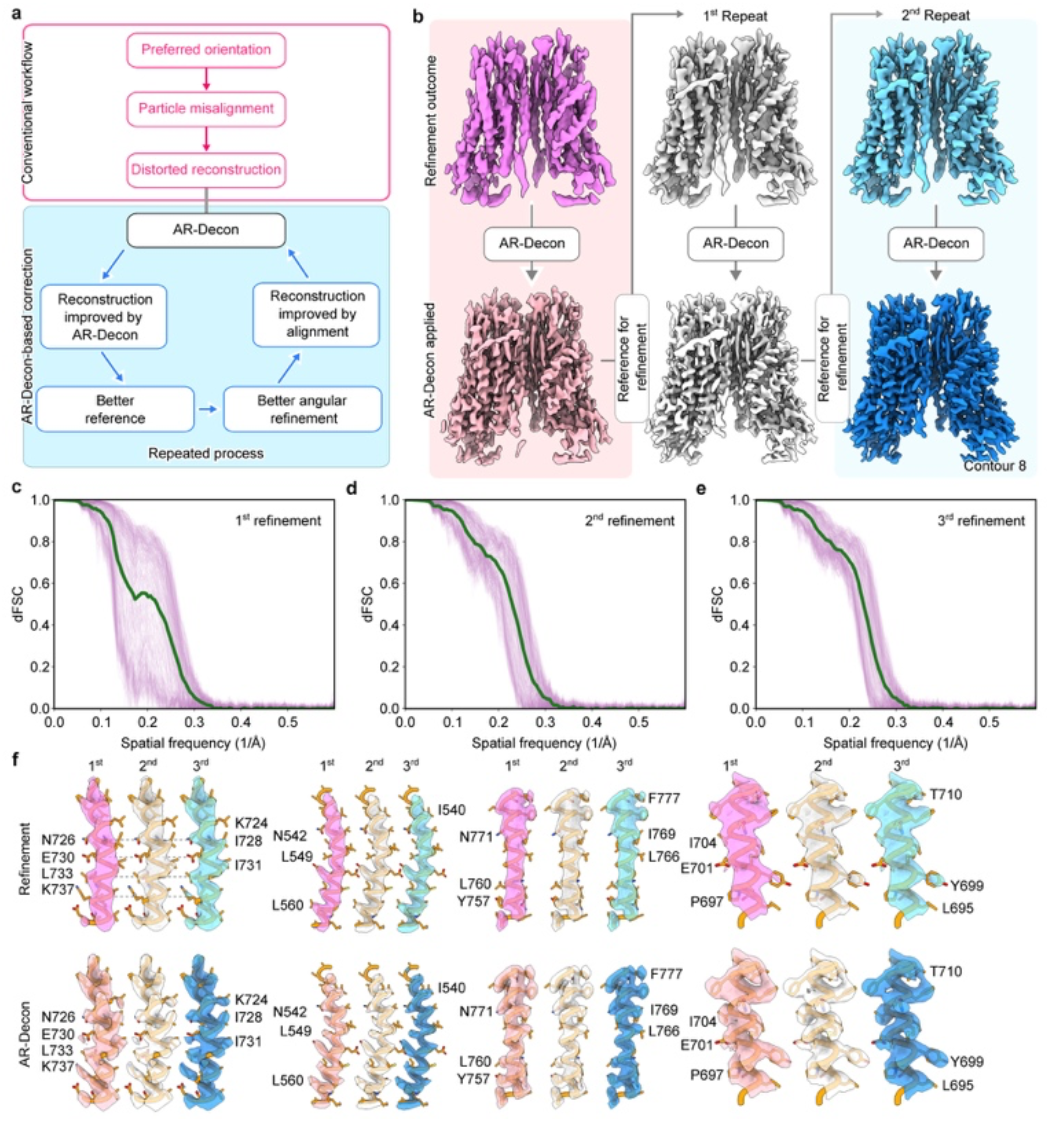
Incorporating deconvolution in map refinement. **a**, Workflow of incorporate AR-Decon with refinement process. **b**, Upper and bottom rows: refined (upper) and deconvolved TMEM16A map (bottom) of each refinement cycle. Left to right: Final map from previous refinement, and two repeated refinements using deconvolved map as the reference map. **c, d, e**, dFSC curves indicate progressive improvement through each round of processing, being calculated without a mask using two half-maps from the initial (**g**), first (**h**) and second (**i**) repeat of refinement. **g**, Detailed comparisons of various helices in the density map after each round of additional refinement.

**Supplementary Figure 5.**
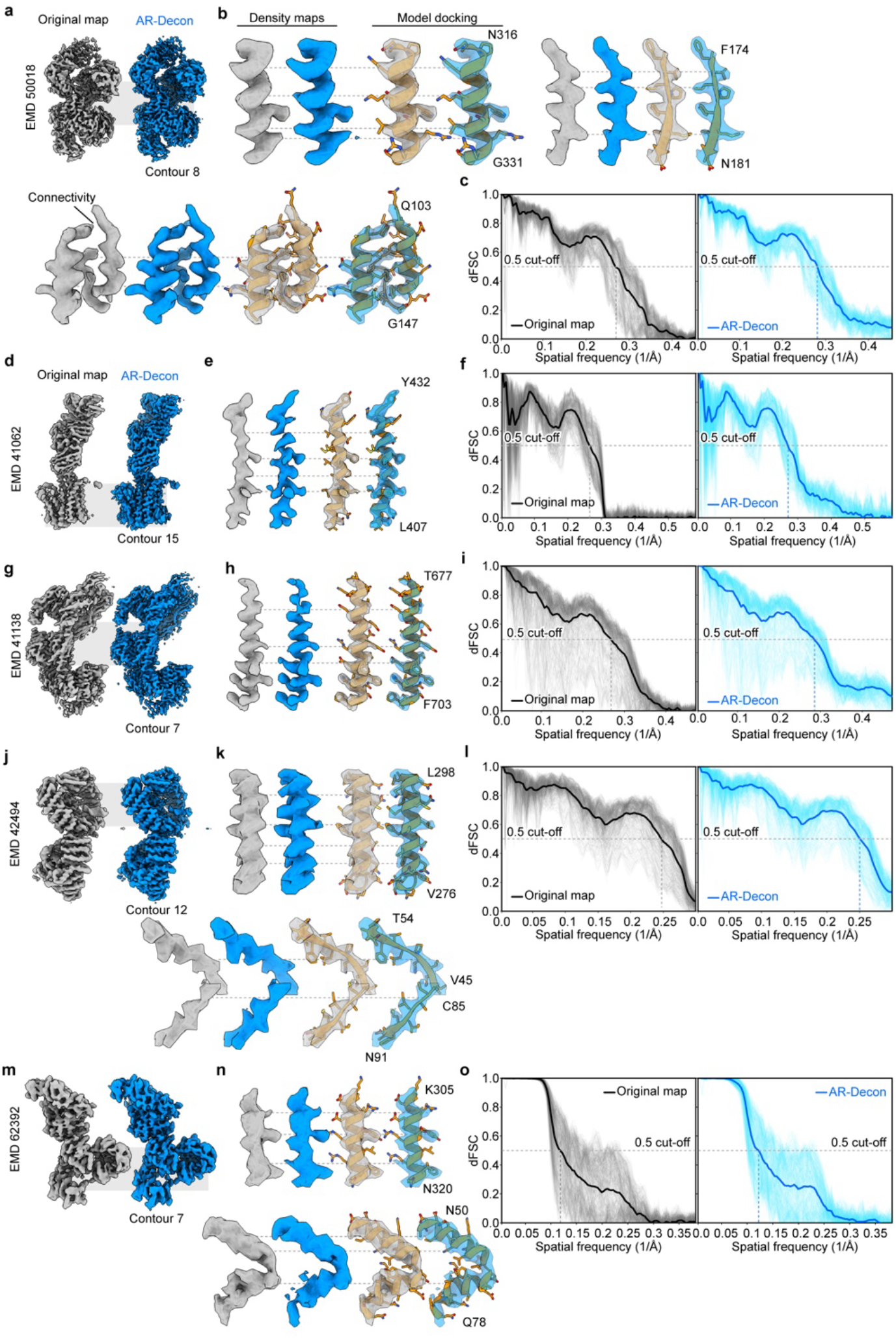
Benchmarking AR-Decon in maps with anisotropic resolution. **a, d, g, j, m**, Global view of EMDB map (grey) and its corresponding deconvolved maps (blue). **b, e, h, k, n**, Selected local regions show improvement of AR-Decon. Densities from the original map are shown in grey, from deconvolved maps are shown in blue. Densities are shown in solid color, or transparent with docket atomic model. **c, f, i, l, o**, Map-to-model dFSC of EMD maps (grey) and deconvolved maps (blue).

**Supplementary Figure 6.**
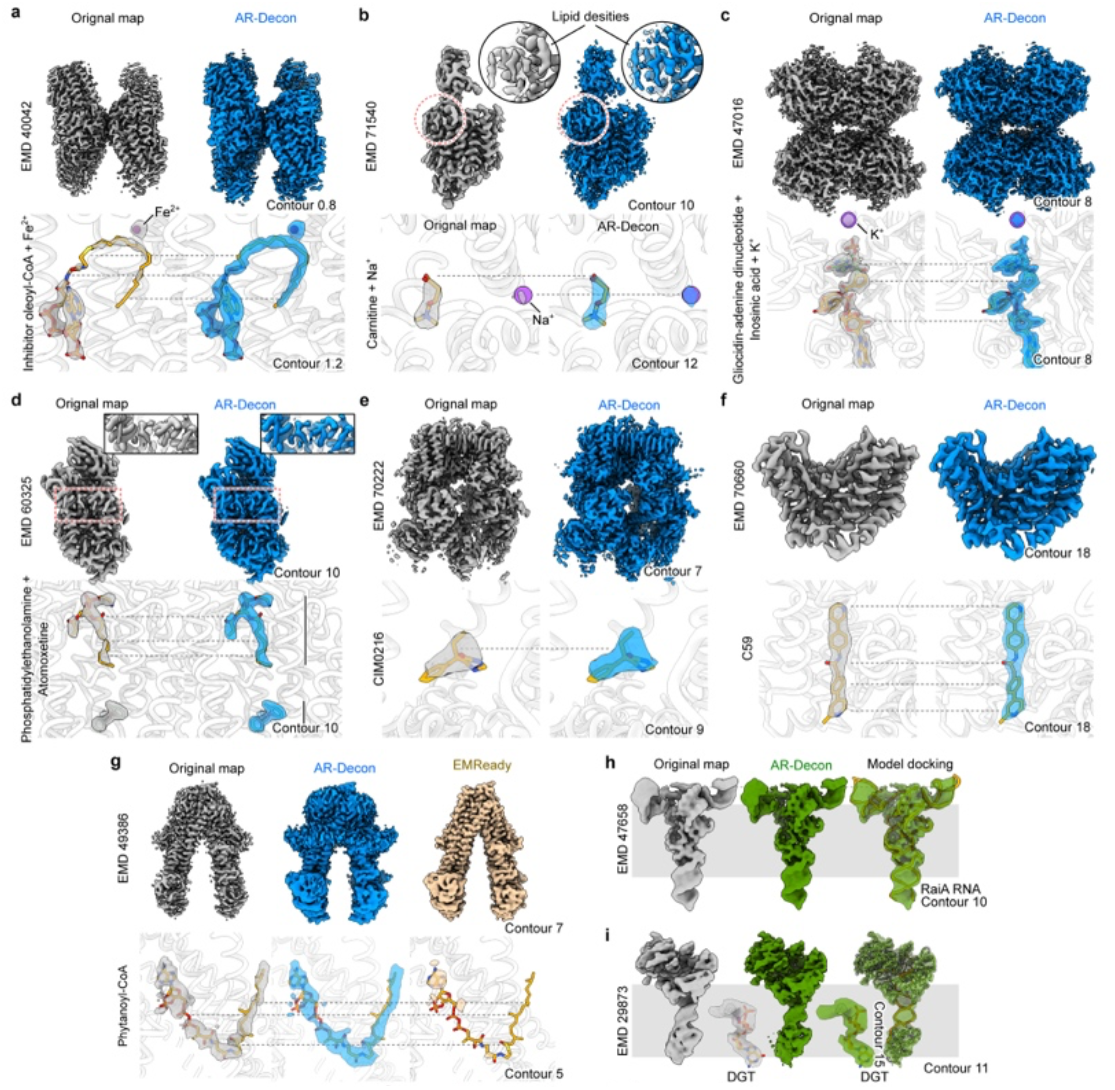
Benchmarking AR-Decon in maps with isotropic resolution. **a** - **f**, upper panel shows the EMDB (grey) and deconvolved (blue) maps. lower panel shows selected non-protein densities from the EMDB (grey) and deconvolved (blue) maps. **g**, Upper panel: original EMDB (grey), deconvolved (blue) and EMReady (yellow) maps of ABCD3 bound with phytanoyl-CoA, shown at the same contour level. Bottom panel: density of phytanoyl-CoA from original (grey), deconvolved (blue) and EMReady (yellow) maps. **h** and **i**, EMDB (grey), AR-Decon (green) maps and deconvolved map with atomic model docked.

**Supplementary Figure 7.**
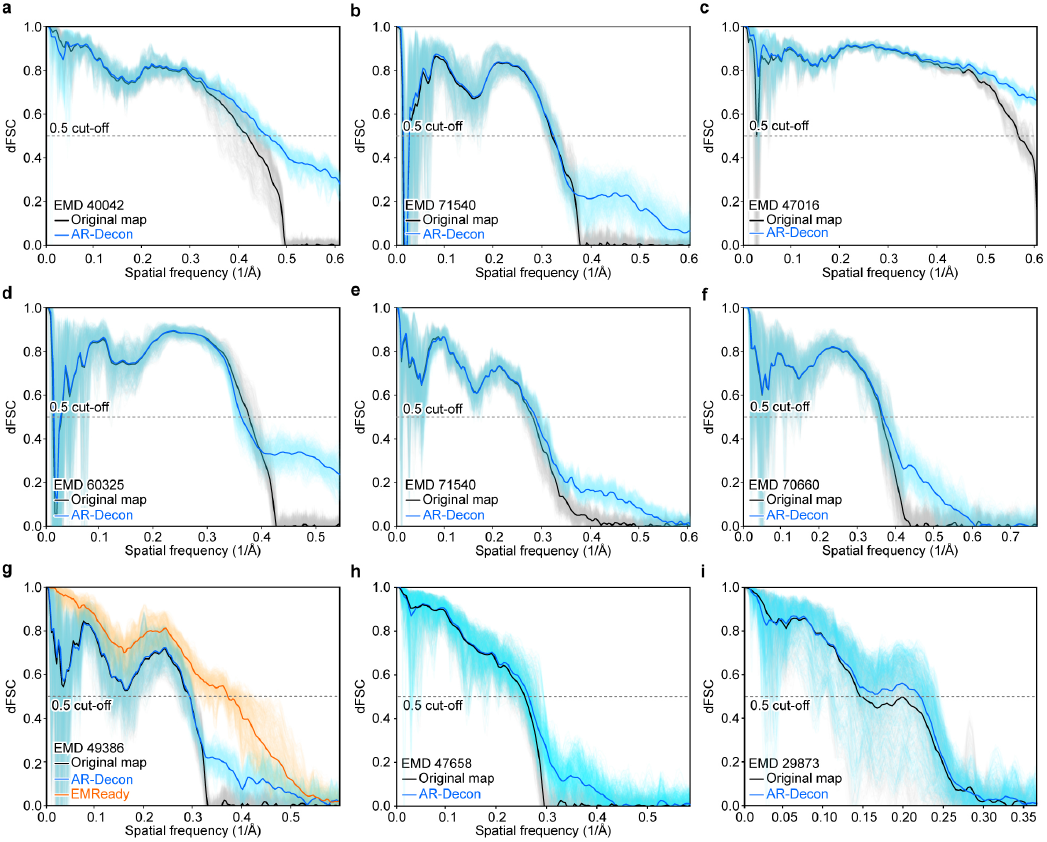
Map-to-model dFSC of benchmarking maps. Map-to-model dFSC of benchmarking maps used in Figure 7a-f.

**Supplementary Figure 8.**
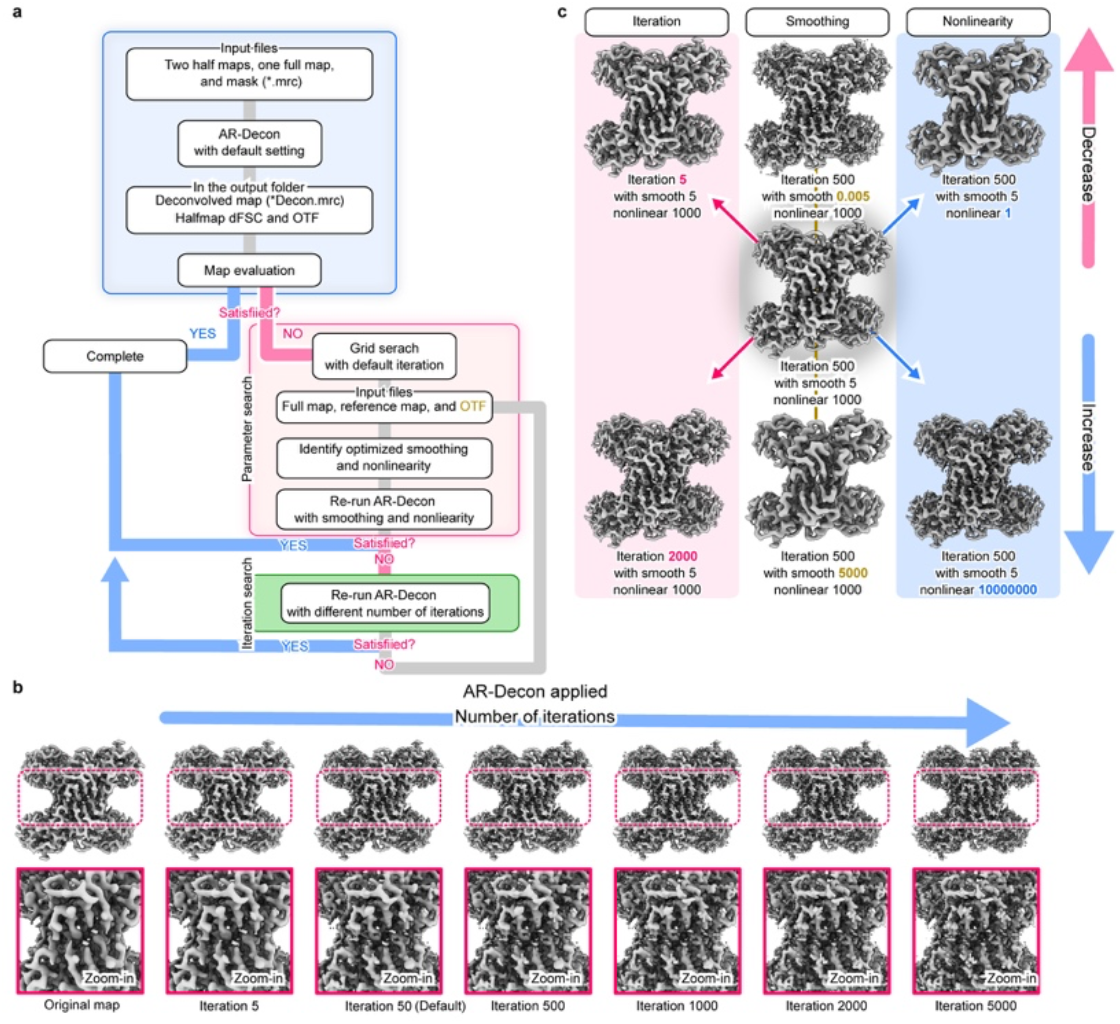
AR-Decon workflow and influence of deconvolution parameters. **a**, Schematic of AR-Decon pipeline. Half-maps, full map and mask are processed with default setting and evaluated. **b**, Deconvolution of EMD-29644 with default regularization parameters but different iteration cycles. **c**, Effect of smoothing and nonlinearity on deconvolution. Parameter setting and iteration numbers are marked.

**Supplementary Figure 9.**
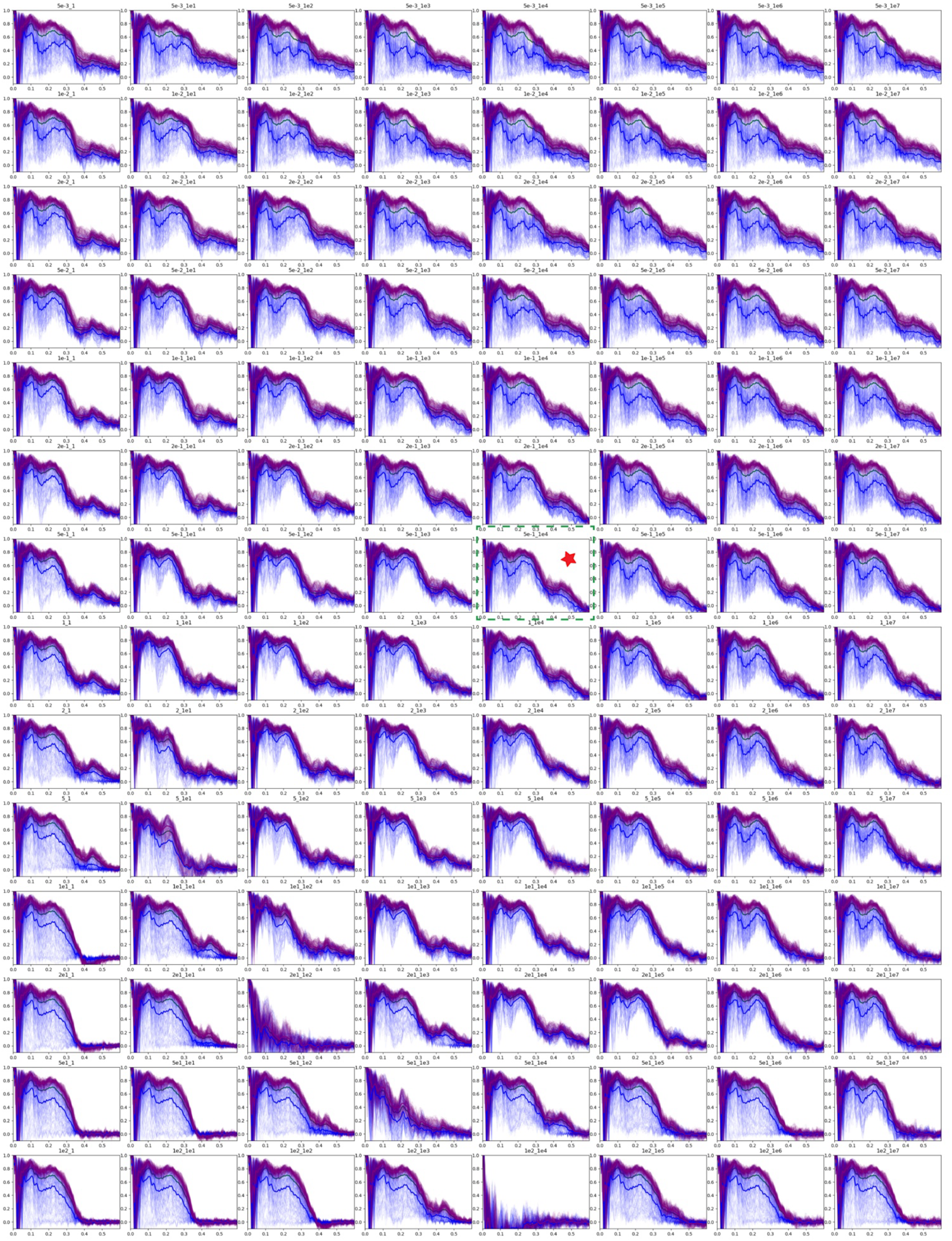
Optimization of regularization parameters for AR-Decon. Gallery of dFSC curves from screening of smoothing and nonlinearity with ferroportin test dataset. Parameters used as default in AR-Decon is marked by a red star.

**Supplementary Figure 10.**
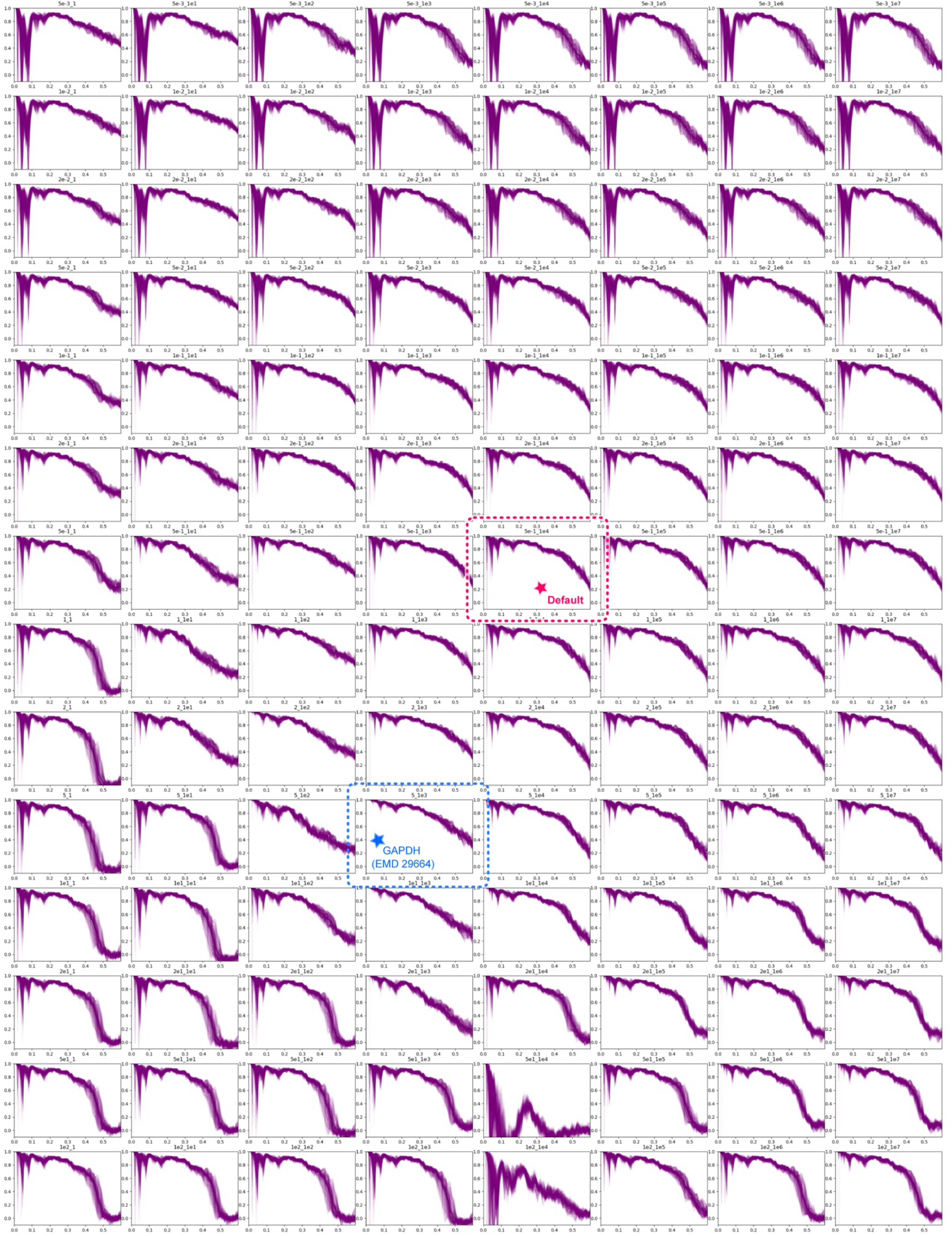
Optimization of regularization parameters with GAPDH. Gallery of dFSC curves from screening smoothing and nonlinearity by deconvolving GAPDH EMD-29664. Parameters marked with star were chosen as a set of regularization parameters for deconvolving EMD-29664 map.

